# NMR Analysis of the interaction of Ethanol With the Nicotinic Acetylcholine Receptor

**DOI:** 10.1101/139717

**Authors:** David Naugler, Robert J. Cushley, Ian Clark-Lewis

**Author notes:** Deceased (Ian Clark-Lewis). **Abbreviations** C_50_: concentration which elicits 50% of effective response EDTA: ethylenediaminetetraacetic acid HEPES: 4-(2-hydroxyethyl)-1-piperazineethanesulfonic acid K_d_: dissociation constant, concentration at which complex is 50% dissociated nAChR: nicotinic acetylcholine receptor NMR: nuclear magnetic resonance NOESY: nuclear Overhauser effect spectroscopy Pi: inorganic phosphate PO_4_^2-^ NLS: nonlinear least squares.

## Abstract

Ethanol exerts its actions in the central and peripheral nervous systems through the direct interactions with several proteins, including ligand-gated ion channels such as the nicotinic acetylcholine receptor (nAChR). The binding interaction between ethanol and sodium cholate solubilized nicotinic acetylcholine receptor protein can be detected through either NMR line broadening or T_1_ titration. In this paper, we examine the use of weighted Navon T_1p_ analysis of T_1_ titration data for the estimation of the dissociation constant of ethanol for the nAChR. We show that Navon T_1p_ analysis underestimates binding affinity. The application of rigorous limits for confidence intervals within a nonlinear regression analysis of this data provides a best estimate of K_d_ = 55 μM at 4 °C. within an unsymmetrical 90% confidence interval of [0.5, 440 μM]. Accordingly, the best estimate of the binding free energy is ΔG^0^, = −5.4 Kcal/mole within a 90% confidence interval of [−8.0, −4.3 Kcal/mole],relative to conventional standard states.

## 1. Introduction

Ethanol can exert profound effects on several components of the central and peripheral nervous systems, including ligand-gated ion channels, [1]. The effects of ethanol can be either stimulatory or inhibitory. For example, ethanol acts as a functional antagonist of ionotropic glutamate receptors [2], but stabilizes the open state of the muscle-type acetylcholine receptor [3]. Extensive studies of the effects of ethanol and other general anesthetics on ligand-gated ion channels provide convincing evidence for the notion that these compounds bind to discrete sites on the channel [4, 5].

The idea that ethanol and other general anesthetics bind to discrete sites on the channel protein has been supported by several site-directed mutagenesis studies on several different ligand-gated ion channels in which specific mutations in the pore-forming domains of the protein were found to alter the sensitivity of the receptors to ethanol [6, 7]. However, these studies should infer that the compound of interest bound to a discrete site on the receptor from looking at effects on functional properties. Due to the low affinity of ethanol (μM to mM range), it is not possible to study the interaction of ethanol with its binding site on the receptor using conventional ligand-binding techniques. However, nuclear magnetic resonance techniques can be used to study the interaction of ethanol with the receptor [8–12], and thus obtain estimates for the true affinity of ethanol for its binding site.

In this study, we have used NMR relaxation methods to estimate the binding parameters for ethanol in the presence of sodium cholate solubilized nicotinic acetylcholine receptor protein. Using this approach, we find that ethanol binds to the solubilized receptor with an affinity in the 100 μM range, and compare this value with estimates of the affinity from measurements that depend on alteration of receptor function. In a previous study [52], we used ^1^H chemical shifts in a nonlinear regression model.

## 2. Materials and methods

### 2.1 Receptor protein isolation and purification

Preparation of sodium cholate (≥99% Sigma)solubilized nicotinic acetylcholine receptor protein generally followed the procedure of ref. [13]. All steps handling the protein were carried out at 4°C, unless otherwise stated. Receptor rich membranes from the electroplax tissue of *Torpedo californica* were prepared from frozen electroplax (Pacific Biomarine, Venice, CA) using the procedure of Sobel et al. [14] as modified by Epstein and Racker [15]. Membranes were stored frozen at −70 ^o^C in 0.4 M sucrose, 2 mM HEPES (≥99.5% Sigma)pH 7.5 until used. Sucrose and HEPES were removed by dialysis against cold distilled water containing ~85 μM EDTA(99.4-100.6% Sigma-Aldrich), which was renewed regularly over several days. Receptor rich membranes were then reduced to pellet form by ultracentrifugation at 130,000 g for 30 min. Pellets were resuspended in ‘Buffer I’, containing 10 mM Na phosphate(96% Sigma), pH 8.0, 100 mM NaCl(≥99.0% Sigma-Aldrich), 60 mM KCl (≥99% Sigma) by sonication. This was followed by ultracentrifugation at 130,000 g for 30 min., to reduce receptor rich membranes to pellet form. A second smaller batch of Buffer I was prepared using D_2_O ((99.9 atom % DAldrich) rather than H_2_O. Using sonication, the receptor rich membranes pellet was resuspended in Buffer I D_2_O solution and the pD adjusted to 10.6 with 1 N NaOD. Alkaline-treated membranes (2.5 mg protein /ml) in Buffer I D_2_O solution were solubilized by the addition of 20% Na cholate/D_2_O to a final cholate concentration of 1%. After incubation for 20-30 min. and ultracentrifugation for 30 min at 130,000 g, the supernatant (cholate extract) was collected and the pellet (membranes) discarded.

Acetylcholine receptor concentration was determined using [^125^I] α-bungarotoxin binding [16].

### 2.2 NMR

All protein NMR was done on a Bruker AMX600 operating at a ^1^H frequency of 600.13 MHz and at a temperature of 277°K. Thermal regulation was achieved using a Bruker Eurotherm Variable Temperature unit. Longitudinal relaxation, T_1_, was studied using a conventional inversion recovery pulse program. T_1_ values were estimated using the nonlinear least squares T_1_/T_2_ routines of Bruker UXNMR or XWinNMR software. Since the T_1_ of free ethanol is 5 sec. at 277 K, this necessitated a recycle time of 25 sec. with 16 scans at each delay. Sixteen variable delays were used. The following list is typical. T_1_ delays (seconds): 25, 16.6, 11.1, 7.4, 4.94, 3.292, 2.194, 1.463, 0.975, 0.650, 0.433, 0.289, 0.193, 0.128, 0.085, and 0.000004. The T_1_ delays used are an important determinant of the precision versus time efficiency of T_1_ estimation. Ref. [17] and references cited therein, provides an analysis for optimal estimation of longitudinal relaxation and advocates the use of unequally spaced sample points in the context of nonlinear least squares parameter estimation. Ref. [18] provides an analysis of the problem of estimating NMR relaxation rate in the context of median estimation [19]. However, it is mathematically more tractable to treat an estimation scheme that is analytical. In ref. [18, 20, 48–50] we develop a forward modeling approach to exponential estimation for the general relaxation model

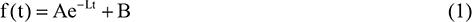

It is shown that under reasonable assumption, the four-point spectral estimator scheme

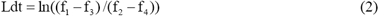

provides optimal estimation. In this scheme, dt is the sample delay between four equally spaced points f_1_, f_2_,f_3_,f_4_. In the context of real error, this scheme is biased, although the bias can be calculated and subtracted off. However, in the context of complex NMR error, spectral estimation [18, 20] is unbiased. We show that to second order, the standard deviation, S_2_, of this estimation scheme is given as,

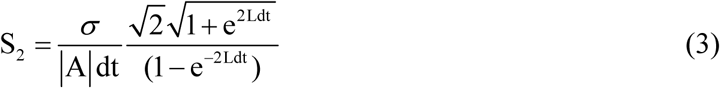

where σ is the noise level.

An analytical, forward modeling approach to an estimation problem is of great value. However, no such approach exists for the estimation problem treated in this paper. The Navon method [21, 22] initially considered here, places special emphasis on the T_1_ value of the free ligand. The T_1_ of ‘free’ ethanol was determined in a sample, which was identical in D_2_O Buffer I and in sodium cholate concentration, to the others except it lacked the 1.7 μM nAChR protein. The ethanol concentration in the ‘free’ sample was 55 mM.

### 2.3 Longitudinal relaxation

Because the transverse relaxation phenomenon is observed on a multiplet, we chose to investigate the concentration dependence of longitudinal relaxation. For two-site fast exchange, nuclear magnetic longitudinal relaxation can be expressed in a mole fraction weighted model [23] as:

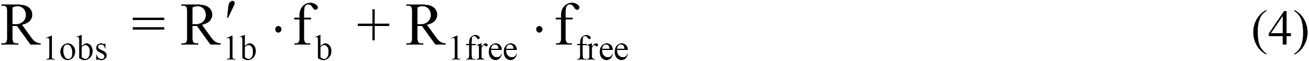

where, R_1obs_, 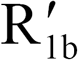 and R_1free_ are the relaxation rates observed and of the bound and free species, respectively, in Hz (1/s). Variables, f_b_ and f_free_are the mole fractions of bound and free species, respectively. These latter quantities are calculated as part of a two-site equilibrium model, which is dependent upon the dissociation constant, K_d_. The relaxation rates of free and bound species can be related to relaxation times [21, 22] by:

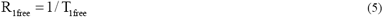

which is a directly measurable quantity, and

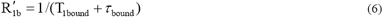

in which T_1bound_ is the relaxation time of the bound species, intrinsic to the nucleus in question, and τ_bound_ is the bound lifetime, intrinsic to the exchange process itself. In ethanol, there are two different ^1^H nuclei detected, the methyl and methylene protons, each with its own R_1obs_, R'_1b_, R_1free_ and T_1bound_. The exchange or bound lifetime is related to kinetic exchange rate by τ_bound_ = 1/ k_-1_.

Since we have a method for an accurate determination of R_1free_, [21, 22], we can rewrite equation (4) as:

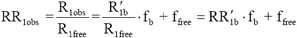

in a reduced rate formulation. This has the advantage of allowing the comparison of like, dimensionless quantities. Table 1 shows the raw data that are transformed. The exchange process can be represented as:

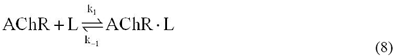

where AChR represents the (nicotinic) acetylcholine receptor, L represents the ligand, ethanol, and AChR · Lrepresents the receptor-ligand complex. Three equations define this two-site exchange system:

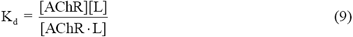

defines the equilibrium condition,

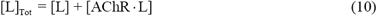

specifies the mass balance in ligand, and

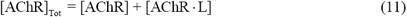

specifies the mass balance in protein.

**Table 1.** T_1_ and T_1p_ Values and Uncertainties

| [EtOH] | $CH_2$ $T_1$ | $CH_3$ $T_1$ | % error | $CH_2$ $T_1$ error | $CH_3$ $T_1$ error | $CH_2$ $T_{1p}$ | $CH_2$ $T_{1p}$ error | $CH_3$ $T_{1p}$ | $CH_3$ $T_{1p}$ error |
| --- | --- | --- | --- | --- | --- | --- | --- | --- | --- |
| 1.34 | 2.949 | 2.394 | 5% | 0.15 | 0.12 | 6.34 | 1.0 | 4.49 | 0.57 |
| 2.67 | 3.904 | 3.121 | 7% | 0.27 | 0.22 | 13.34 | 4.51 | 7.96 | 1.88 |
| 3.89 | 4.577 | 3.942 | 8% | 0.37 | 0.32 | 26.86 | 18.23 | 16.98 | 8.15 |
| 7.23 | 5.059 | 4.254 | 9% | 0.46 | 0.38 | 60.94 | 94.65 | 24.82 | 17.59 |
| free | 5.517 | 5.134 | 10% | 0.55 | 0.51 |  |  |  |  |

We have a system of three equations in three unknowns that can be solved, in principle. Because we do not initially know the value of K_d_, we can start with the fiction that its value is zero. This is a tight binding approximation. Since the ligand is in excess, we then have

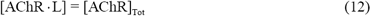

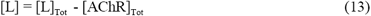

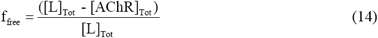

and

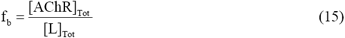

for substitution into equation (7). This gives us a model that we can regress against the raw data given in Table 1,

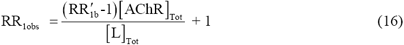

According to Navon analysis, error in Riot translates into error in the determination of (T_1_b + τb) but not in Kd. The (T_1_b + τb) values found here are like those of Navon [21]. Therefore, any errors or uncertainties in the value of [AChR]_TOT_ will have little or no effect on the results.

Figure 1 shows the line broadening of ethanol that occurs in the presence of nAChR protein. The multiplet structure of ethanol is completely washed out due to a line broadening of 20 Hz. The ^3^J_HH_ coupling constant of 7 Hz gives rise to a triplet at the methyl and a quartet at the methylene, when this can be resolved. The line broadening declines as the cholate detergent denatures the protein. Storage at 4 °C will not prevent complete denaturation over a period of a month. After denaturation, the multiplet structure of ethanol then is observed again (data not shown).

**Figure 1.**
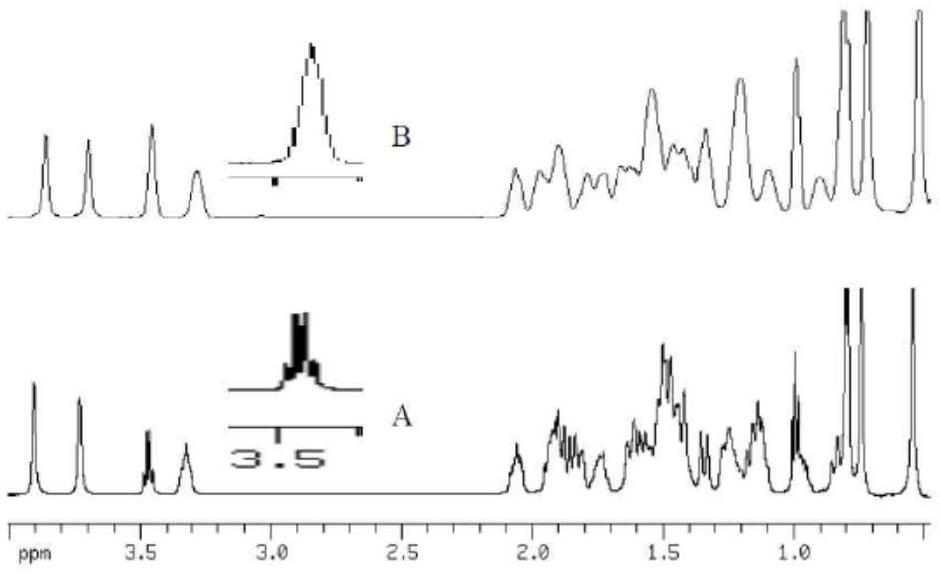
The bottom trace, A, shows the spectrum of 1% sodium cholate in D2O with ethanol at approximately 7 mM. In the spectrum of the top trace, B, nAC'hR protein (1.7 |uM, not visible) is solubilized with 1% sodium cholate in D2O and ethanol is present at approximately 7 mM. The resonance at 1.0 ppm is the methyl and that at 3.5 ppm is the methylene of ethanol.

The large value for the correlation of determination for the tight binding model means that the estimation of K_d_ is a difficult task. The Navon method [22, 24] provides one approach. This method is an approximate linearization that is analogous to some methods of linearization of hyperbolic data in Michaelis-Menton kinetics. It relies on plotting T_1p_, defined by,

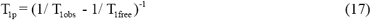

versus ligand concentration. Historically, the notation above was used because of the practicalities of experimental physics. An equivalent notation would have,

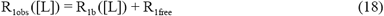

Conceptually, this is very compelling because with effort, it is possible to obtain a very good estimate for T_1free_. In this approximation, K_d_ is determined as the negative x-intercept.

However, the estimation of errors in T_1_ is troublesome. For instance, Mao [25] performed a study of 200 inversion-recovery experiments on ^17^O relaxation in water. He found that the statistical error in T_1_ is about six times as large as that reported by nonlinear least squares software. NLS software usually assumes linear approximation inference regions [26], and this assumption is often found to be inadequate. We advocate implementing an exact forward modeling approach to the problem of estimating mono-exponential decay in NMR.

The results of fitting the data to equation (16) are displayed in Figure 2. This is linear regression because the unknown parameters in equation (7) appear in that equation as linear parameters, although the calculated fitting functions are curved. There is great advantage to linear regression. For one, the statistical estimates of the parameters are well behaved. The coefficient of determination in this analysis is 0.9686. This analysis leads to values for (T_1bound_ + τ_bound_) of 0.0054 ± 0.0008 s for CH_3_ and 0.0074 ± 0.0011 s for CH_2_. Since these values are significantly different (p < 0.1), it can be concluded that differences in T_1bound_ for ethanol CH_2_ and CH_3_ account for the difference and that τ_bound_ is small by comparison, otherwise the ratio of T_1bound_ for CH_2_ and CH_3_ would be unreasonable. If τ_bound_ is small then this is one more piece of supporting evidence that this chemical exchange occurs in the fast exchange limit. The values for ethanol CH_2_ and CH_3_ here are of similar magnitude to the value of (T_1bound_ + τ_bound_) = (9.9 ± 0.8) X 10x^3^ s, for nicotine binding to asolectin solubilized nAChR found by authors in ref. [21], by monitoring relaxation of the proton *para* to the aromatic nitrogen. This similarity is supporting evidence for a likeness in stoichiometry of binding [18, 21].

**Figure 2.**
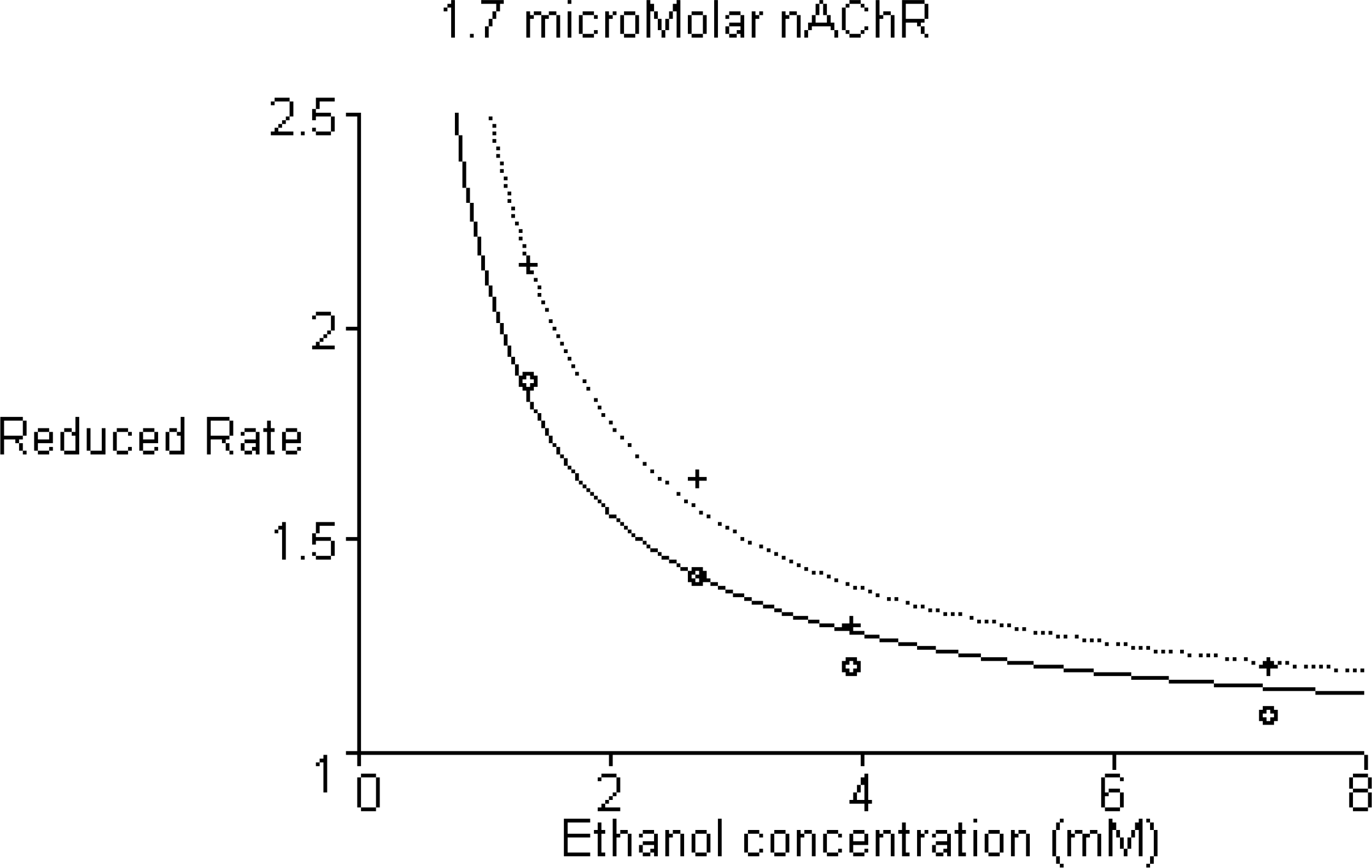
Plot of reduced relaxation rate of ethanol CH_3_ and CH_2_ protons versus ethanol concentration. A least square fit of the tight binding model, Kd=0 reproduces the relaxation data with coefficient of determination 0.976 The model reproduces relaxation data for ‘free’ ethanol exactly. The solid line and circles relate to CH_2_ protons, dotted line and crosses relate to CH_3_ protons.

Given that τ_bound_ ≪:T_1bound_ then τ_bound_ < 0.0027 ± 0.0004 s, and k_-1_ >370 + 55s^−1^.

The residuals displayed in Figure 2 provide insight into the nature of the error. It is seen that there is a consistency in absolute error in R_1_. Consequently, the relative error in R_1_ is inversely related to the magnitude of R_1_ and proportional to T_1_. As the magnitude of T_1_ grows, its absolute error grows even faster. This is consistent with experience. Long T_1_ times are difficult to measure accurately. Tables I and II show the calculation of the propagation of errors as percentage and absolute errors in T_1_ for each concentration and thence to T_1p_ errors and weights. Figure 3 displays a T_1p_ transformation of the data and curves of Figure 2, which suggests the use of extrapolation for the determination of K_d_. Extrapolation of points at low concentration provides an estimate of K_d_= 122 ± 258 μM. (See discussion below.) The T_1_ values and uncertainties estimated in Table 1 are used to calculate T_1p_ values and weights in Table 2. Also, shown in Table 2 are fitted values and residuals according to the model:

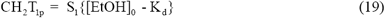

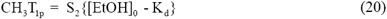

and shown as Navon analysis in Figure 3. In this linear analysis, the intercept has a large standard deviation 0.188 and a small positive value 0.0188. Accordingly, we set the best estimate, 0 < K_d_ < 0.17 mM. From this analysis [27], we can set a 95% confidence range for K_d_ as [0, 0.46 mM].

**Table 2.** T_1p_ Values, Weights and Fit

| [EtOH] | $CH_2$ $T_{1p}$ | $CH_2$ weight | $CH_3$ $T_{1p}$ | $CH_3$ weight | $CH_2$ $T_{1p}$ calculated | $CH_3$ $T_{1p}$ calculated | $CH_2$ $T_{1p}$ residuals | $CH_3$ $T_{1p}$ residuals |
| --- | --- | --- | --- | --- | --- | --- | --- | --- |
| 1.34 | 6.34 | 1 | 4.49 | 3.077 | 6.462 | 4.395 | -0.122 | 0.0949 |
| 2.67 | 13.34 | 0.049 | 7.96 | 0.28 | 12.967 | 8.819 | 0.372 | -0.859 |
| 3.89 | 26.86 | 0.0030 | 16.98 | 0.015 | 18.934 | 12.877 | 7.925 | 4.102 |
| 7.23 | 60.94 | 0.00011 | 24.82 | 0.0032 | 35.271 | 23.988 | 25.668 | 0.831 |

**Figure 3.**
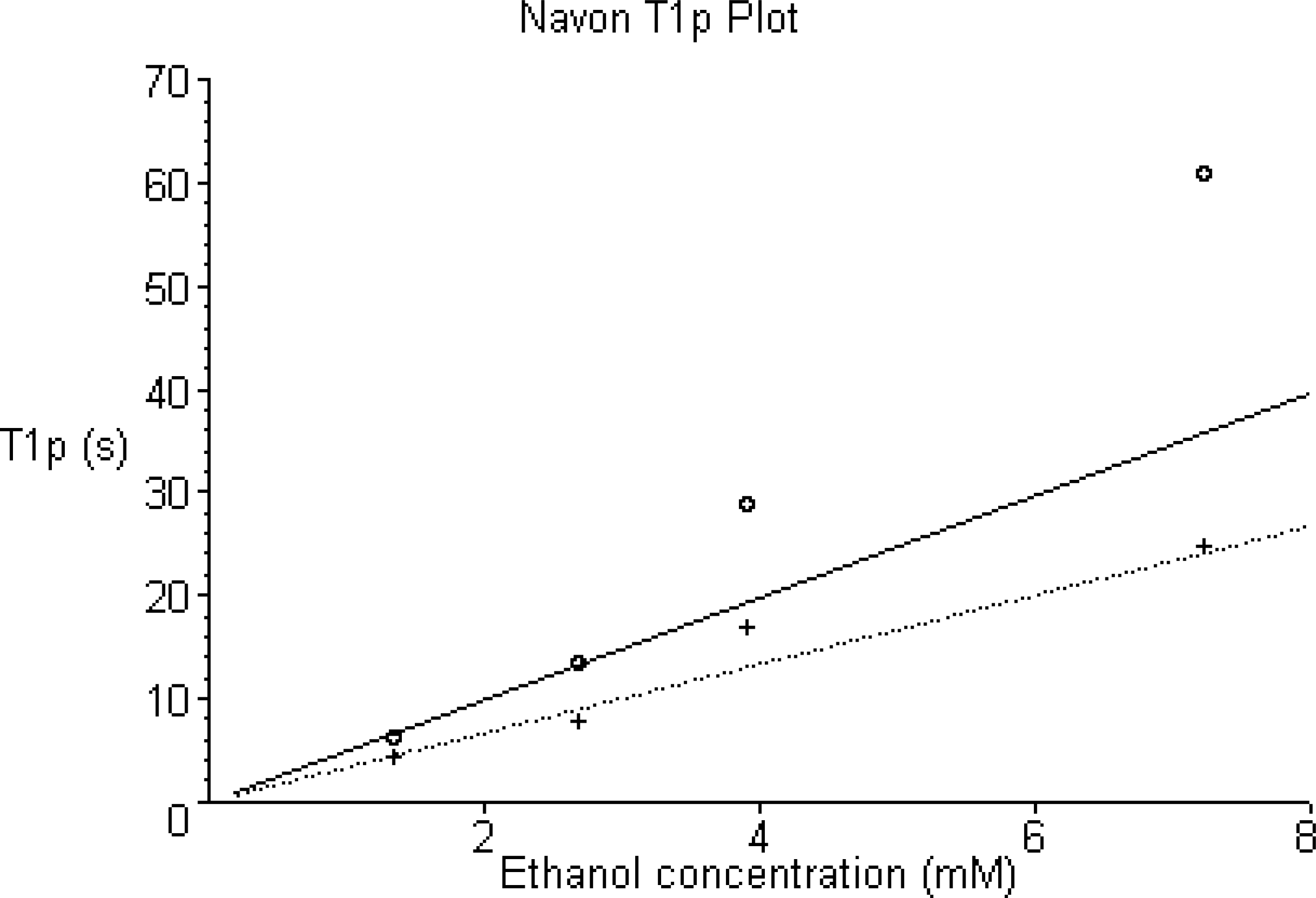
Navon T_1p_ plot of CH_3_ and CH_2_ proton relaxation data versus ethanol concentration shows how the data and least squares curves of Figure 2 transform under the T _1p_ formulation. The solid line and circles relate to CH_2_ protons, dotted line and crosses relate to CH_3_ protons. The T_1p_ value of ‘free’ ethanol is infinite because of numerical instability. Extrapolation of points at low concentration provides an estimate of ^K_d_^ = 122 + 258 μM.

#### Evaluation of Navon Analysis

Navon analysis [21, 22] is an approximation which provides approximate conclusions but which avoids otherwise very difficult analysis. The level of approximation involved in the Navon treatment may be justified when the level of approximation involved in the treatment of NMR relaxation is considered. However, since we propose an exact treatment of NMR relaxation, a second look at the Navon approximation is warranted.

The three equations that define equilibrium in a two-site chemical exchange can be solved [18]. To simplify notation, we use E to represent AChR. These equations are:

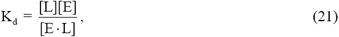

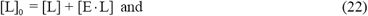

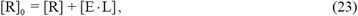

a system of three equations in three unknowns that is solved when:

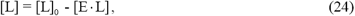

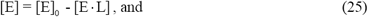

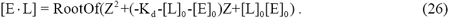

We must choose the positive root of the polynomial for a physically reasonable solution. Hence,

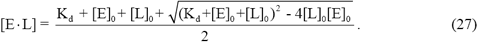

[L]_0_ is the independent variable, the initial ligand concentration. Using this exact formulation of equilibrium, we can simulate [18] the functional form of NMR relaxation with assumed parameter values. For example, Figure 4 plots the functional form of T_1obs_ for two-site exchange given some presumed values. Examination of the shape of the curve at low ligand concentration shows that this curve is not exactly rectangular hyperbolic, which is assumed for Navon linearization. Figure 5 shows the exact functional form of T_1p_ given some presumed parameter values. It is not exactly a straight line, curvature at low concentrations is pronounced. In this simulation, a value of K_d_ = 0.1 mM was assumed but extrapolation of the straight portion of the curve back unto the negative x-axis predicts a value of K_d_ = 0.2 mM, according to the Navon method. Hence, we can say that the Navon method is likely to underestimate binding affinity by a factor of about two.

**Figure 4.**
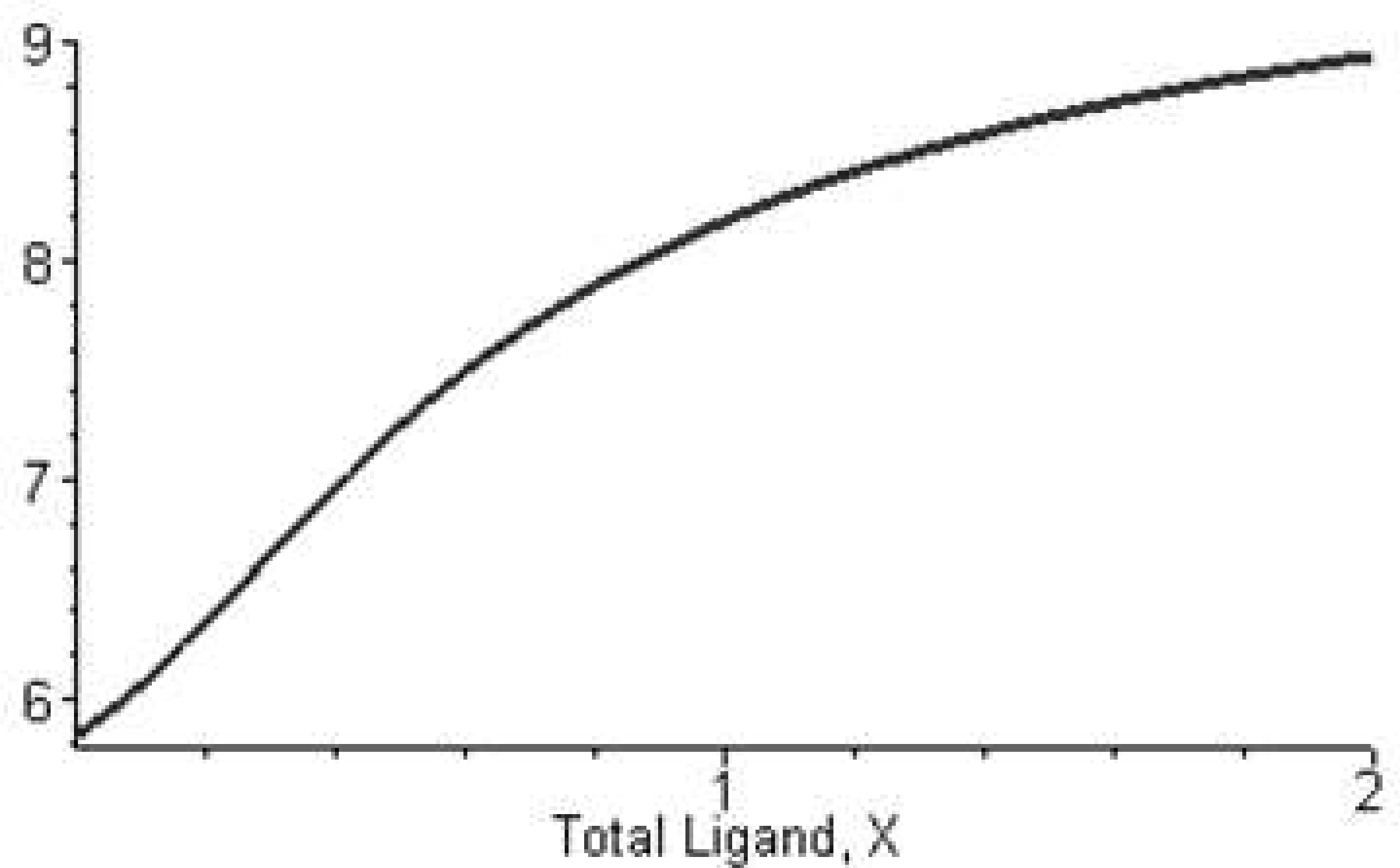
Plot of predicted T_1_ versus ethanol concentration, assuming K_d_ = 0.1 mM, showing the calculated variation of T_1_ with variation of total ligand concentration, X = [L]_0_.

**Figure 5.**
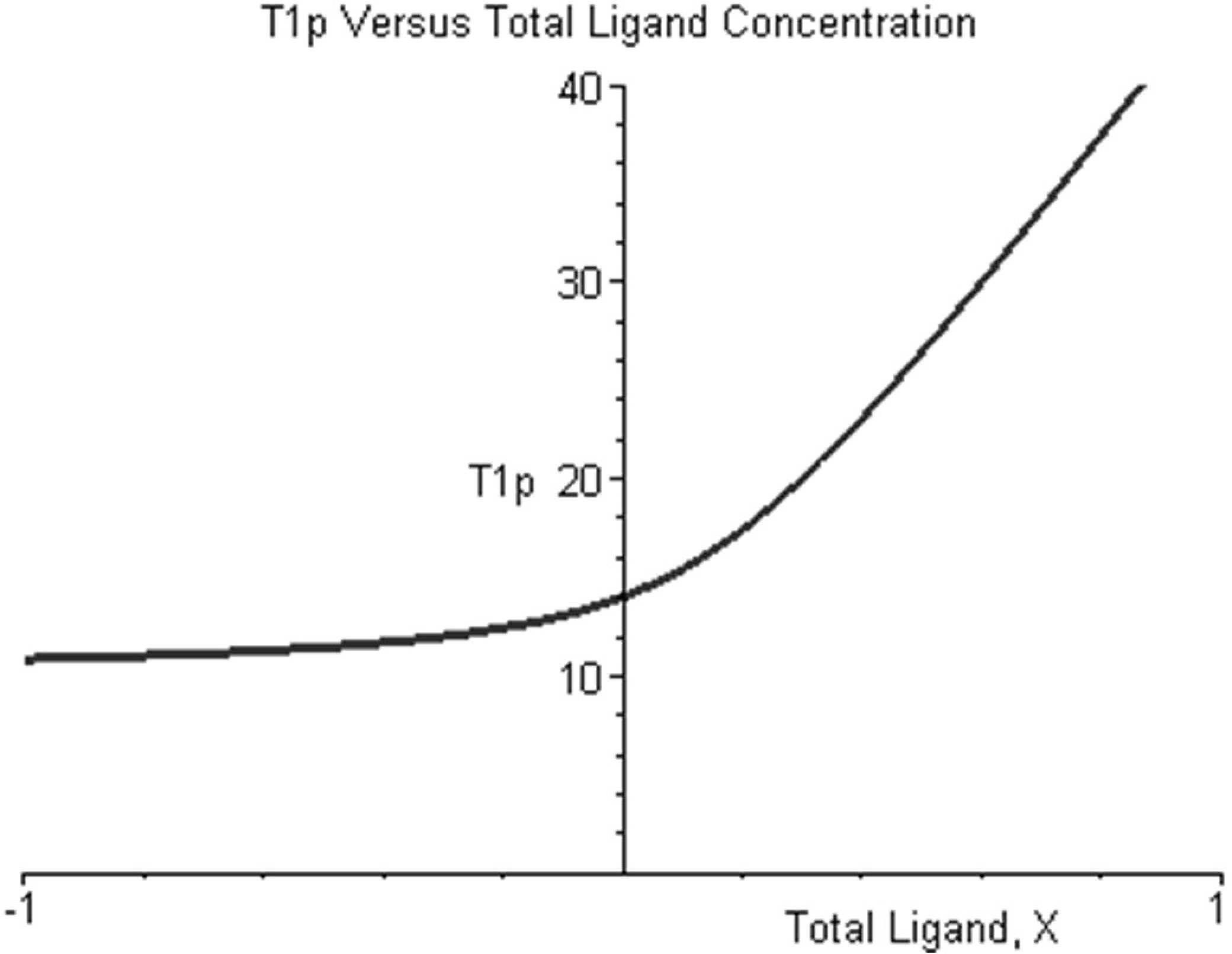
Plot of the calculated variation of T_1_p with variation of total ligand concentration and setting K_d_ = 0.1 mM. For purposes of demonstration, the plot is extended into the region of physically unrealizable negative ligand concentration. On the right, the curve is asymptotically linear. Extrapolation from the linear region onto the negative x-axis gives an intercept of −0.2 mM.

We had said that weighted Navon analysis predicts that 0 < K_d_ < 170 μM for a 95% confidence interval of [0, 460 μM], for the binding of ethanol to nAChR. With this new evaluation of the Navon method, we are better to say that analysis of the data predicts that 0 < K_d_ < 85 μM. This value is consistent with some estimates of the affinity of ethanol for neuronal AChRs [28]. Extrapolation of points at low concentration would suggest K_d_ is around 61 μM. Additional analysis [18] in the context of rigorous limits [27, 29] for nonlinear regression analysis that does not assume linear approximation inference regions [26], provides a best estimate of K_d_ = 55 μM within a 90% confidence range of [0.5, 440 μM]. This similarity to the 95% confidence interval for Navon analysis supports the use of the Navon approximation.

#### A Putative Site of Action

The ethanol molecule can be considered as a rigid electric dipole. When two isolated dipoles achieve a position of closet approach, energy minimization selects an orientation in which the two dipoles cancel, creating effectively, a nonpolar assembly. Sacco and Holtz [10,11] used NMR measurement of intermolecular dipole-dipole relaxation rates and of self–diffusion coefficients of ethanol molecules to establish that ethanol self-association occurs in aqueous media, but only at molar concentrations. An ethanol dimer retains an electric quadrupole moment that is effective over short range. Cohen et al. [30] used [^3^H]acetylcholine mustard to identify residues contributing to the cation-binding subsite of the nicotinic acetylcholine receptor. A representative sequence alignment of the cation-binding subsite is shown below:

~~~
LPSDDVWLPDLVLYNNADGDFAIVHMTK *Torpedo* californica (Pacific electric ray)
VPSEMIWIPDIVLYNNADGEFAVTHMTK alpha polypeptide 2 (human neuronal)
IPSELIWRPDIVLYNNADGDFAVTHLTK alpha polypeptide 4 (human neuronal)
IPSEKIWRPDLVLYNNADGDFAIVKFTK alpha polypeptide 1 (human neuromuscular)
~~~

BLAST search [31] of the nonredundant protein database at GenBank [32] reveals that the sequence PDIVLYNNADG, centered about tyrosine-93 (human neuromuscular numbering), is conserved in fish, mammal, bird, insect and amphibian. The importance of the cation-binding domain is further illustrated in the human neuromuscular junction where it is found to be adjacent to the Main Immunogenic Region (MIR), α67-76 of the autoimmune disease Myasthenia Gravis [33].

We studied [18] the interaction of ethanol and the 28-residue cation-binding subsite, α80-107, of human neuromuscular nAChR. Secondary structure prediction algorithms [34–40] assign very weak structure forming elements to this sequence [34–40]. This prediction is consistent with observations made using heteronuclear and multidimensional NMR methods [18]. There is little consensus among the various secondary structure prediction algorithms, except for a prediction of beta-sheet near the central tyrosine. Examination of the Kyte-Doolittle hydropathy plot [41] for this sequence shows a relatively flat hydropathy in the middle of the peptide, with a more hydrophilic N-terminal end and more hydrophobic C-terminal end. If a 28 residue peptide sequence is truly a random coil, it is predicted to sample nearly 9^27^ = 4.39E695 backbone conformations. Given the shape of the hydropathy plot, we predict this peptide to be condensed in aqueous solution and to sample a smaller conformation space.

The formation of nonpolar molecular assemblies is disfavored in a polar medium but favored in a nonpolar environment. Energy minimization predicts that if an ethanol dimer is found within the nonpolar environment of a protein near an ionizable phenolic tyrosine hydroxyl, that the proton and two ethanol molecules should form a charged nonpolar molecular assembly. Furthermore, such an assembly is predicted to have about the same volume and charge distribution as the choline head group. We observed [18] dramatic change in the proton spectrum of 28mer, most notably in the amide region, induced in the peptide by choline, indication of a change in the ensemble of sampled conformations.

NMR relaxation studies [18] of the interaction of ethanol and cation-binding subsite peptide of human neuromuscular nAChR established that ethanol binds with a stoichiometry of two. Downfield chemical shifts of methylene protons, like a model system [18], indicate that bound ethanol interacts with a proton. From nonlinear regression analysis, the best estimate of one binding constant (that of least affinity) is K_d_ = 39 ± 13 μM at 4 °C. This estimate is within the uncertainty range of the best estimate of the binding affinity of ethanol for the nicotinic acetylcholine receptor, K_d_ = 55 μM. This establishes that the cationic-binding subsite is a putative site of action of ethanol and that tyrosine-93 is implicated in the binding.

## Discussion

In this study, we have used nuclear magnetic resonance techniques to examine the interaction of ethanol with the acetylcholine receptor. Studies on the effects of ethanol on ligand-gated ion channels such as the acetylcholine receptor have used various functional measurements to study the interaction of ethanol with the receptor [51]. While these types of studies have provided great insight into the mechanism of ethanol’s effects on the channel, and even the site(s) of action, they do not directly study the binding reaction between ethanol and the receptor. NMR relaxation methods offer one means of directly measuring the affinity of ethanol for the receptor.

The first biological NMR binding constant determination used chemical shift measurements to study the association equilibria of N-acetyl-D-glucosamine anomers and lysozyme [42]. Later, the determination of binding constants by line width measurements was applied to sulphanilamide binding to bovine carbonic anhydrase [43]. This work was followed by the determination of *τ*_c_ for various inhibitors bound to carbonic anhydrase, using T_1_ and T_2_ measurements at three magnetic field strengths [44]. Biological NMR binding constant determination in the context of nicotinic pharmacology, was first applied to the binding of nicotine to asolectin solubilized nAChR protein using the selective T_1p_ method [21]. This work was followed by binding constant measurement for various ligands to recombinant active site peptides of the nicotinic acetylcholine receptor using the selective T_1p_ method [22]. NMR binding constant estimates agree with estimates obtained by other methods (when available) to within experimental error. A recent validation of this statement, compared NMR binding measurements to kinetics binding measurements, for Vanadium(V) binding to ribonuclease A [45]. This work was later confirmed by other authors [46].

In this study, we obtained an estimate for the dissociation constant for ethanol from cholate-solubilized nicotinic acetylcholine receptors on the order of 50-100 μM. This value is like the concentrations of ethanol that altered the gating of AChRs in PC12 cells [28]. However, this is much lower than blood alcohol levels associated with intoxication (on the order of 20 mM), as well as the concentrations of ethanol that enhance *Torpedo* nAChR gating [47].

The discrepancy between the estimate of the dissociation constant for ethanol binding to the Torpedo receptor and functional measurements on the same receptor could be due to several possibilities. First, the receptor in the NMR experiments may be in the denatured state, and thus ethanol is binding to a completely different conformation of the receptor than studies in the functional state. We consider this unlikely because of the line-broadening of ethanol, which disappears with time as the protein denatures. Since this line broadening is observed in the time scale of our experiments, we assume that the protein is still in a native-like conformation. Second, and more likely, is that like all direct measurements of binding, the site with the highest affinity is the easiest to detect, and in the case of the Torpedo AChR, the highest-affinity site is not necessarily the one responsible for most of the functional consequences of the interaction of ethanol with the receptor. If this is true, then while we have determined the affinity of a site for the interaction of ethanol with the AChR, it may not be the one associated with the alterations in nAChR gating.

Even if the site we have studied is not the one responsible for functional alterations associated with ethanol’s actions, we have demonstrated that it is possible to use NMR analysis to examine the interaction of small-molecules with ligand-gated ion channels. Extension of this technique to any one of several channel modulators with affinities in the micromolar range (which are generally too low to measure with conventional ligand-binding techniques) should allow one to begin to understand the nature of the interaction of these modulators with the channels, and potentially help determine the site(s) of action of these compounds.

## Acknowledgments

The authors are pleased to thank Dr. Michael M. White, Department of Pharmacology and Physiology, Drexel University College of Medicine, for preparation of receptor rich membranes from the electroplax tissue of Torpedo *californica,* and Dr. Ian Clark-Lewis, Biochemistry and Molecular Biology, University of British Columbia, for synthesis of the cation binding-site peptide.

